## Supplementary materials for "WiPP: Workflow for improved Peak Picking for Gas Chromatography-Mass Spectrometry (GC-MS) data"

### Supplementary data

#### Sample preparation (Datasets number 1 and 2)

*Substance mixtures*

Substance mixtures were prepared as described in the method [25]. Briefly stock mixes of compounds 1mg/mL in 20% methanol were combined to give high concentration sample 1:1. This and aqueous dilutions (1:10, 1:100) were extracted as described [25] with modifications. Samples were re-dissolved in water and 2 mL of methanol:chloroform:water (5:2:1), containing 2 µg/mL cinnamic acid per millilitre of (diluted) standard mix were added. Samples were shaken for 30 min at 4 °C to ensure phase equilibration, centrifuged and 550 µL of polar phase were aliquoted and dried for 4 h at 25 °C in a temperature controlled rotational vacuum concentrator (Labconco).

Derivatization: Derivatization was carried out as described [25]. In short, the dried samples were re-dissolved in pyridine (Carl Roth) containing 40 mg $mL^{-1}$ methoxyamine hydrochloride (Sigma-Aldrich) and shaken for 90 min at 30 °C. Afterwards, MSTFA (N-methy-N-(trimethylsilyl) trifluoroacetamide, Macherey-Nagel GmbH & Co. KG, Düren, Germany) for silylation containing nine *n*-alkanes ($C_{10}$, $C_{12}$, $C_{15}$, $C_{17}$, $C_{19}$, $C_{22}$, $C_{28}$, $C_{32}$, $C_{36}$, 25 µg/mL) as retention index markers was added, samples were incubated for 60 min at 37° C under constant shaking. They were centrifuged, and supernatants transferred to glass vials.

*Gas chromatography*

Gas chromatographic separation of compounds was performed as previously described [25] on an Agilent 6890N (Agilent, Santa Clara, CA, USA) equipped with a VF-5ms column of 30 m length (Varian, Palo Alto, CA, USA). The initial temperature of 67.5 °C was held constant for 2 min, before heating with a temperature gradient of 5 °C $min^{-1}$ until 120 °C, followed by a gradient of 7 °C $min^{-1}$ until 200 °C, followed by the third final gradient of 12 °C $min^{-1}$ until 320 °C where the temperature was then held at 320 °C for 6 min. The transfer line was kept at 250 °C throughout. A cold injection system was used with a matching baffled deactivated liner (CIS4, Gerstel, Mülheim an der Ruhr, Germany), operating in split mode (split 1:5, injection volume 1 µL), with the following temperature gradient applied: hold of the initial temperature of 80 °C for 0.25 min, followed by a temperature increase of 12 °C $s^{-1}$ to 120 °C, followed by a temperature increase of 7 °C $s^{-1}$ to 300 °C with a hold time of 2 min.

*Sequence setup*

Samples were measured in 10 blocks of decreasing dilution (1:100, 1:10, 1:1) with 2 washes (containing only MSTFA and retention index standards) in between to counteract possible carryover.

*Low resolution dataset*

Derivatization and gas chromatographic separation were carried out as described above. MS measurement was performed on a Pegasus 4D-TOF-MS-System (LECO Corp., St. Joseph, MI, USA) with 1 Da mass resolution and −70 eV electron impact ionization and acquisition voltage of 1700 V complemented with an auto-sampler (MultiPurpose Sampler 2 XL, Gerstel, Mülheim an der Ruhr, Germany). Spectra were recorded in a mass range of 60 *m*/*z* – 600 *m*/*z* with an acquisition rate of 10 scans/s.

*High resolution dataset*

Derivatization and gas chromatographic separation were done as described above. MS measurement was performed on a 7200 Q-TOF (Agilent, Santa Clara, CA, USA) with a mass accuracy of 5 ppm, using −70 eV EI. Spectra were recorded in a mass range of 60 *m*/*z* –600 *m*/*z* with an acquisition rate of 10 scans/s.

**Library matching**

Detected peaks were matched against an in-house library based on the cosine similarity of normalized spectra and the RI difference. The similarity was reduced by a factor corresponding with RI deviation (0.97, 0.95, 0.93, 0.85 for RI differences larger than 1.5, 3, 4, 5, respectively), scores above 0.9 were considered as matches.

**Peak set scoring**

To score quality and quantity of detected peaks, both assigned peak classes and number of peaks within each class are considered. Therefore, a class-score is assigned to each of the seven peak classes according to their quality. Correctly detected peaks are rewarded with positive class-scores, detected peaks which are poor quality are penalized with negative class-scores. The overall score for all detected peaks is the sum of the number of peaks within each class multiplied by the corresponding class-score. The scoring function can be written as:

$$score = \sum_{i=1}^{7} n_i \cdot r_i$$

Where $i$ is the class, $n$ the number of detected peaks within this class and $r$ the reward/penalty value for $i$. Class-scores can be defined in the config file> "optimisation-score", or default values are retained. The class-scores balance the coverage of compound-related peaks with the detection of poor quality or noise peaks. Naturally, the highest quality, true positive peaks are scored highest (default: 2) and apex shifted ones second highest (default: 1). All other classes should be scored negatively, as they represent poor quality peaks (default: -2). Wrongly detected peak borders indicate unsuitable parameters while true peaks which are incorrectly classified as noise indicate algorithmic difficulties on the data. Thus, exact class-scores are context dependent and might vary with instrumentation, sample and biological question.

**Peak sampling for training data generation**

Peak picking algorithms with a range of parameters are run on all training samples. This produces a pool of training peaks which can be used for manual annotation. The peak-pool contains duplicates as different parameters might result in the selection of the same peak. To minimize the number of duplicates, peaks within the pool were merged based on their retention time in three subsequent steps: First, peaks with the same apex but slightly varying borders; second, peaks sharing the same borders but a slightly varying apex; third, peaks with slightly varying borders and a slightly varying apex. The threshold for the merging steps is user defined and can be adjusted in the config file> merging (default: 0.2).

Subsequently, a user defined number $n$ of peaks (default: 200; config file> training_data-general) is sampled from the merged-peak-pool. To ensure that peaks are sampled from the whole retention time range, peaks are sorted by retention time and split into 10 equally sized groups. From within each group, $\frac{n}{10}$ peaks are sampled uniformly without replacement. Remaining peaks (if any) are sampled uniformly without replacement independent of the retention time. If the pool contains less than $n$ peaks, all peaks are plotted.

**Peak processing for SVM classification**

For peak classification via SVM, the original peak data (a N x M intensity matrix, where N is the number of recorded m/z values (EICs) and M the number of recorded spectra within the peaks RT) was processed by the following steps:

1. Linear interpolation in the RT dimension.

This is necessary because machine learning classification algorithms require input data with a fixed shape, but detected peaks vary in width and therefore in shape (the M dimension of the intensity matrix). To affect as few peaks as possible, the average peak width of the training data set was used as reference width. After this step, the intensity matrix of all peaks is of shape N x O, where O is the number of spectra corresponding with the average peak width.

2. Normalization to [0, 1]. By normalization, low intensity peaks do not differ from high intensity peaks anymore, thus, only peak shape and not absolute intensity values are considered.

3. Reordering of the rows (EICs) of the intensity matrix in descending order, based on the maximum normalized intensity value within each row. This step avoids training on particular rows (m/z values).

4. Conversion of the processed intensity matrix to an intensity vector. Machine learning classification algorithms require a vector as an input, therefore, the rows of the intensity matrix are placed into a single row. The intensity vector is of length N*O.

After processing, a peak is represented by a vector of size N*O with values between 0 and 1. These vectors can be placed in a N*O dimensional space.

During training, a SVM segments this space into X compartments, where X is the number of predefined classes. The segmentation is performed in a way that best fits the classified training data. To classify a new vector, it is simply placed in the N*O dimensional space and the corresponding compartment is the predicted class.

### Supplementary figures

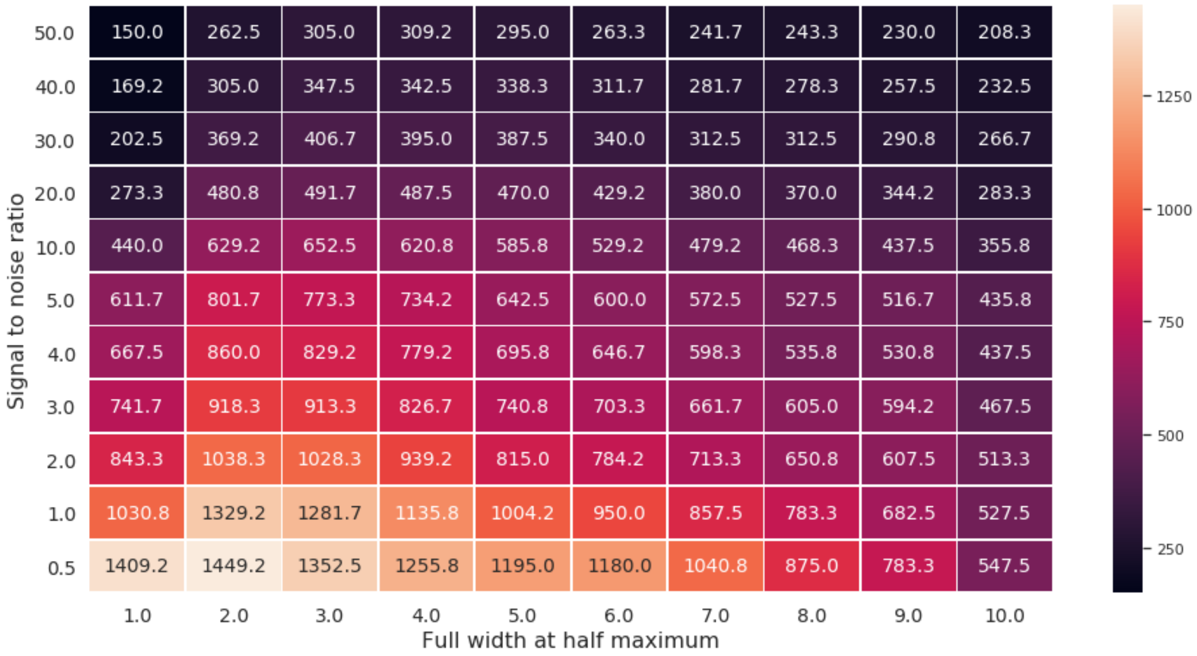


**Figure S1.** Parameter optimisation results. Each of the cells correspond to a parameter set for the matchFilter algorithm and contain the computed scores (See scoring function in supplementary methods). The optimal parameters for this dataset were assessed to be i) Full width at half maximum height (FWHM) = 2 ii) signal to noise ratio = 0.5. For clarity purposes, only two parameter optimisations are shown here. The grid search may cover more parameter dimensions when three or more parameters are optimised.

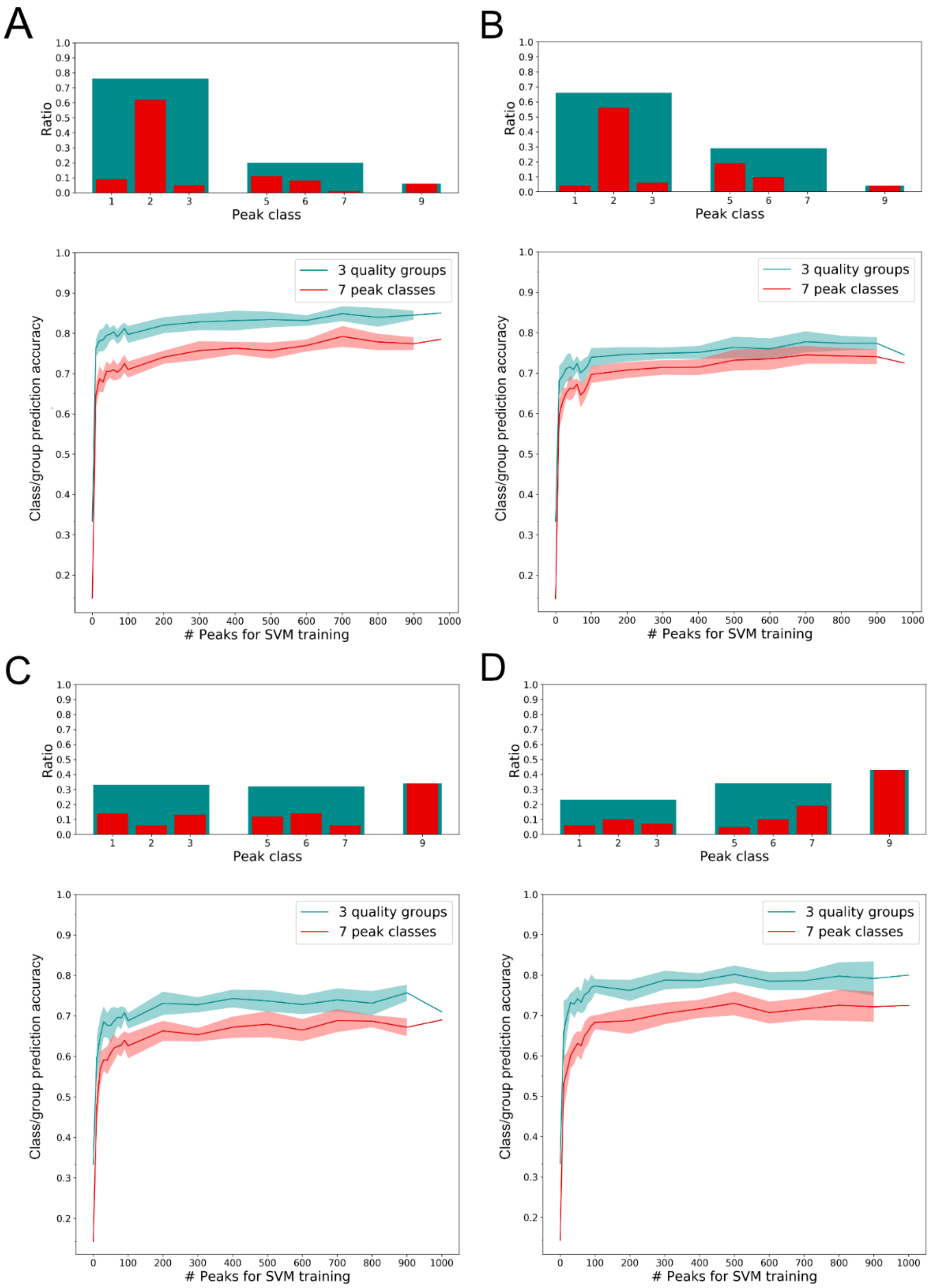


**Figure S2.** Bar charts represent the distribution of classes of 1000 different peaks as annotated by an individual experienced in mass spectrometry. The graph shows the class prediction accuracy as a function of the training size. Red denotes the individual 7 peak classes as defined in WiPP, Green denotes the further classification of these into the 3 measure-of-quality classes used for the scoring function (high quality, intermediate quality and noise peaks). **A** & **B**: Low resolution data (Dataset 1). **C** & **D**: High resolution data (Dataset 2). **A** & **C**: XCMS MatchedFilter. **B** & **D**: XCMS centWave. In the graph, the differences between the two curves is due to intra group misassignment. The peaks used for training were subsampled according to the class distribution shown in the barplot. Peak classes: 1 – Apex shifted to the left. 2 – Centred apex. 3 – Apex shifted to the right. 5 – Merged/shoulder peak. 6 – Peak with wide margins to window border. 7 – Peak exceeds window border. 9 - Noise

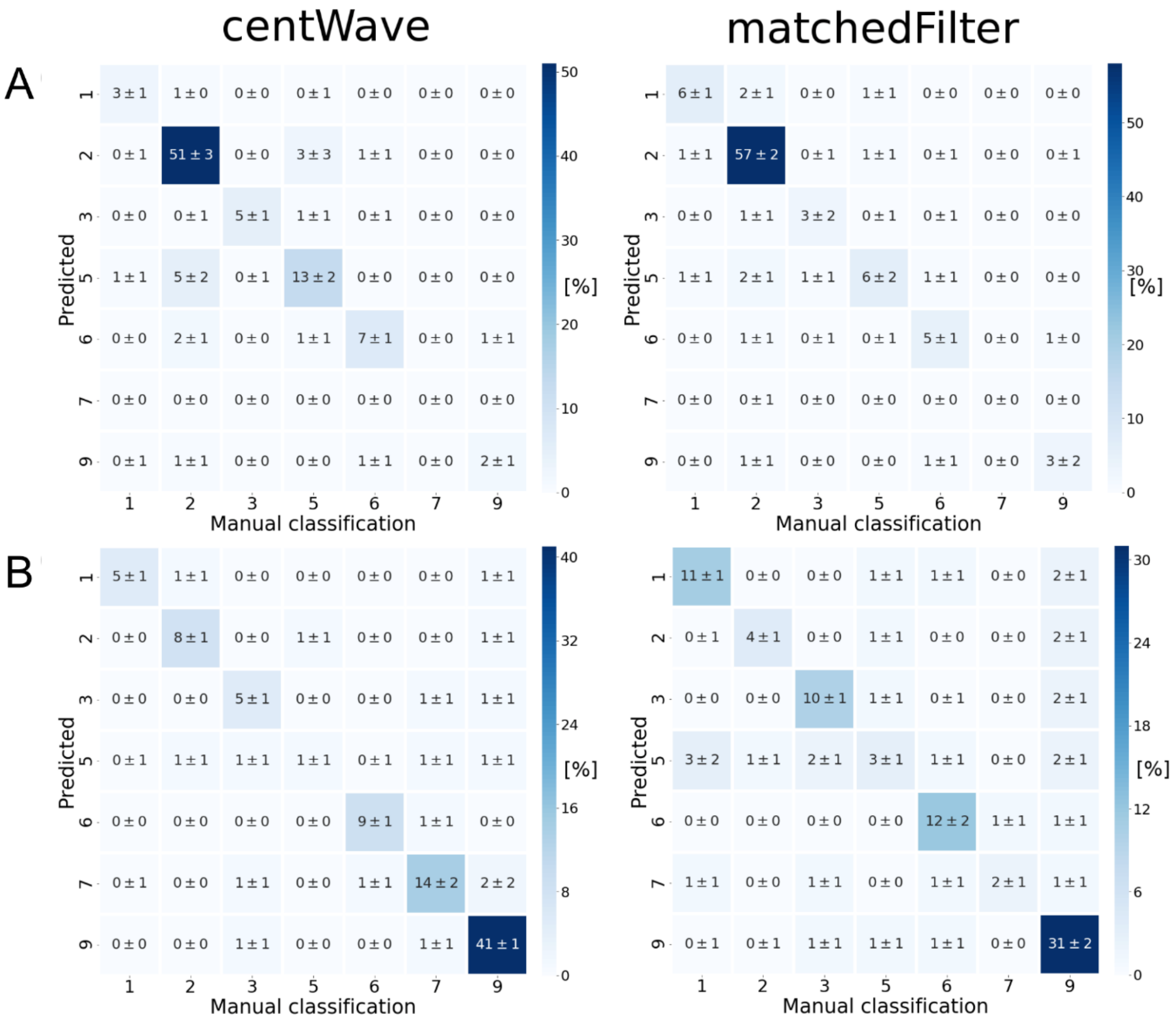


**Figure S3.** Confusion matrices comparing manually assigned classes with classes predicted by trained SVM classifiers. **A**: Low resolution data (Dataset 1). **B**: High resolution data (Dataset 2). The prediction was performed using a stratified five-fold cross-validation for SVM training.

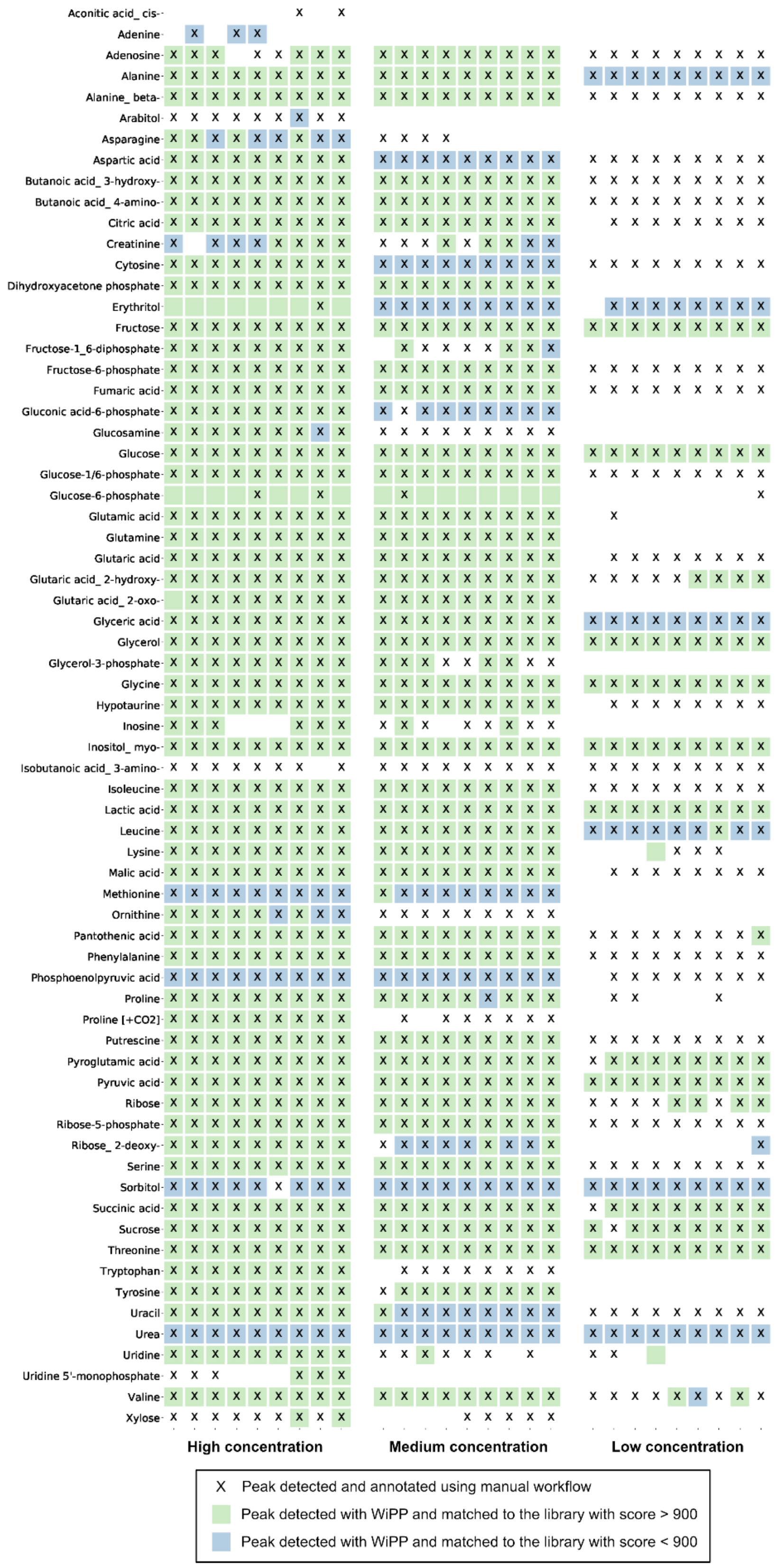


**Figure S4.** Comparison of the list of metabolites detected and annotated manually or automatically by WiPP in the three concentration of the dataset 1.

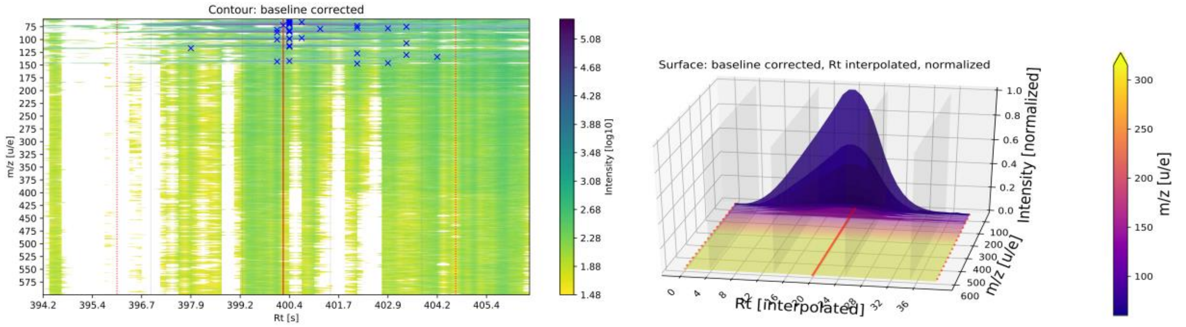


**Figure S5.** WiPP peak visualisation. **A**. Baseline corrected surface plot showing the boundaries of the peak detected by the peak picking algorithm as dotted red lines and the detected apex as a solid red line. x and y axes represent retention time and *m*/*z* respectively. The blue crosses represent the highest intensity for individual *m*/*z* values within the retention time window, a user defined intensity threshold allows the display of blue crosses. **B**. Baseline corrected, interpolated and normalized 3-dimensional plot showing the peak detected by the peak picking algorithm. Red dotted and solid lines indicate peak boundaries and apex respectively.

### Supplementary table

**Table S1.** Optimal algorithm parameters found for the two datasets using WiPP and IPO * High resolution data only.

| | **Dataset 1** | | **Dataset 2** | | |
|---|---|---|---|---|---|
| | **centWave** | **matchedFilter WiPP** | **centWave WiPP** | **matchedFilter WiPP** | **matchedFilter IPO** |
| **pwMin** | 2 | NA | 2 | NA | NA |
| **pwMax** | 3 | NA | 6 | NA | NA |
| **fwhm** | NA | 3 | NA | 1 | 8.8 |
| **sn** | NA | 0.5 | NA | 10 | 10.1 |
| ***ppm** | NA | NA | 5 | NA | NA |
| ***step** | NA | NA | NA | 0.1 | 0.05 |
| ***steps** | NA | NA | NA | 1 | 1 |
| ***mzdiff** | NA | NA | 0.2 | -0.5 | 0.75 |

**Table S2.** Parameter set and ranges used to generate the training data. * High resolution data only.

| **XCMS centWave** | | **XCMS matchedFilter** | |
|---|---|---|---|
| **pwMin** | 1, 2.5, 5 | **fwhm** | 2.5, 5, 7.5 |
| **pwMax** | 5, 10, 15 | **sn** | 1, 2.5, 5 |
| ***mzdiff** | -0.1, 0, 0.1, 0.5 | ***mzdiff** | -0.1, 0, 0.1, 0.5 |
| **ppm** | 5, 10, 20 | ***step** | 0.1, 0.25, 0.5 |
| | | ***steps** | 1, 2, 3 |

**Table S3.** Chromatof pre-processing parameters for manual annotation of Dataset 1.

| Chromatof parameter | Value |
|---|---|
| Data reduction rate | 4 |
| Cut mass range | 70 - 600 |
| Baseline offset | 1 (just above the noise) |
| Number of points for smoothing | Auto |
| Max number of peaks | 600 |
| S/N | 20 |
